# Mouse population genetics phenocopies heterogeneity of human *Chd8* haploinsufficiency

**DOI:** 10.1101/2022.08.26.504147

**Authors:** Manal Tabbaa, Allison Knoll, Pat Levitt

## Abstract

Preclinical models of neurodevelopmental disorders typically use single inbred strains which fail to capture human genetic and symptom heterogeneity that is common clinically. We tested if systematically modeling human genetic diversity in mouse genetic reference panels would recapitulate population and individual differences in responses to a syndromic mutation in the high-confidence autism risk gene, *CHD8*. Trait disruptions mimicked those seen in human populations, including high penetrance of macrocephaly and disrupted behavior, but with robust strain and sex differences. For every trait, some strains exhibited a range of large effect size disruptions, sometimes in opposite directions, and remarkably others expressed resilience. Thus, systematically introducing genetic diversity into mouse models of neurodevelopmental disorders provides a better framework for discovering individual differences in symptom etiologies and improved treatments.

**One-Sentence Summary:** Autism trait heterogeneity due to a syndromic gene mutation is recapitulated in mice by incorporating genetic diversity.

## Main Text

Individuals exhibit striking clinical heterogeneity in the presence and severity of neurodevelopmental disorder (NDD) symptoms and co-occurring behaviors, even with the same rare and highly penetrant gene mutation (1,2). The predominant use of single isogenic strains in preclinical *in vivo* models of NDDs has prevailed for decades, but this *‘n of 1’* genome strategy can be improved upon by using genetic approaches that capture the biobehavioral and genetic heterogeneity observed in patient populations (3–6). Fields that have modeled human genetic diversity by using panels of genetically diverse recombinant inbred mouse strains have fueled mechanistic discoveries of individual differences in disorder risk for infectious disease, cancer, immunology, metabolic and neurodegenerative disorders, among others (7–13). In contrast, the contribution of genetic background to the etiology and pathophysiology of NDDs is far less understood, presenting significant barriers towards developing robust translational models needed for treatment discoveries. Data-rich multi-strain studies that incorporate broad and reproducible variation in genetic background (*i*.*e*., *strain)* can reveal genetic and environmental modifiers of single-gene effects on behavioral outcomes and facilitate i) discovery of etiological mechanisms underlying symptom differences, ii) predictions of which individuals are most at risk, and iii) opportunities to leverage specific backgrinounds for mechanistic based treatment discoveries. Using a mouse genetic reference panel (GRP) approach, we have previously demonstrated strain heterogeneity and heritability of complex traits in wild-type (WT) mice including fear learning and affiliative social behavior (14,15). Here, we report the results of a systematic analysis of the penetrance of trait disruptions caused by a high-confidence autism spectrum disorder (ASD) risk gene across diverse genomes using two GRPs, the Collaborative Cross (CC) and BXD collections that are derived from fully-sequenced founder strains (16–18). Into each GRP strain, we introduced clinically relevant haploinsufficiency of chromodomain helicase DNA-binding protein 8 (*Chd8)* followed by comprehensive phenotyping. *CHD8* encodes a protein that regulates chromatin remodeling and gene expression during brain development (19). In clinical populations, functional mutations in *CHD8* are associated with macrocephaly, ASD, intellectual disability, and anxiety with variable penetrance and severity (20–24).

B6-*Chd8*^*+/-*^ dams were mated with sires from 27 CC, 5 BXD, and C57BL/6J (B6) strains. Maternal genotype was held constant, and sires were removed prior to litter birth to control for strain differences in parental care. The resulting F1 WT and *Chd8*^*+/-*^ male (n = 521) and female (n = 520) littermates across 33 strains were comprehensively characterized (**Fig. 1**), with a focus on phenotypes relevant to *CHD8* haploinsufficiency clinical symptoms.

**Figure 1.**
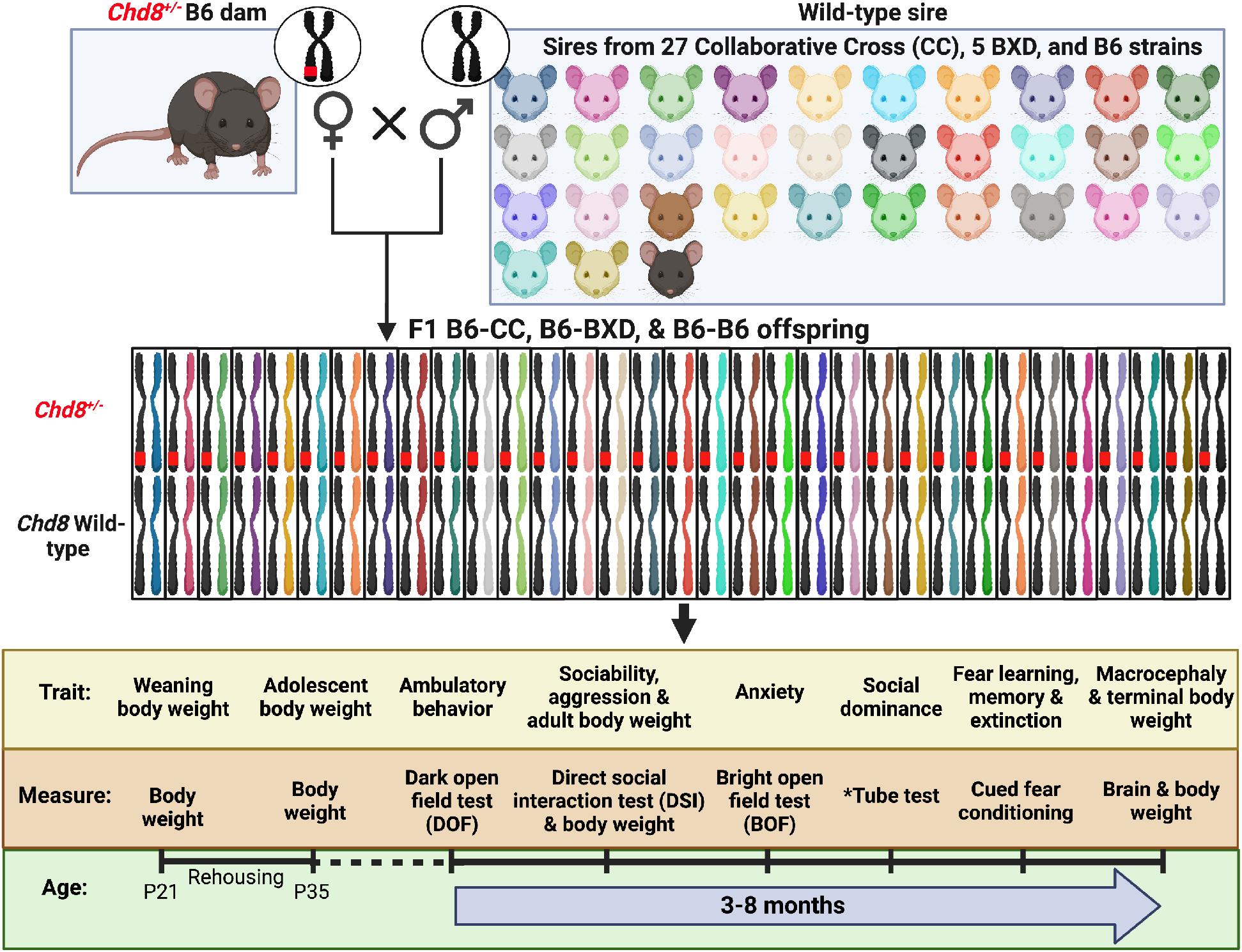
Study Design: Modeling population and individual differences in phenotypic responses to *Chd8* haploinsufficiency. *Chd8* heterozygous (*Chd8*^*+/-*^) B6 dams were mated with wild-type (WT) sires from 27 Collaborative Cross (CC), 5 BXD, and B6 strains to produce F1 B6-CC, B6-BXD, and B6-B6 male and female WT and *Chd8*^*+/-*^ offspring. Subjects were weaned at postnatal day (P) 21 and rehoused prior to P35 to 2 WT and 2 *Chd8*^*+/-*^ same -sex and -strain mice per cage, with littermates preferentially housed together. Behavioral testing began at a mean (∼) age of P115 and was conducted in the order shown on the timeline. Weaning, adolescent (P35), adult (∼P125), and terminal (∼P220) body weights, in addition to terminal brain weights, were measured. 33 strains (8 subjects per *Chd8* genotype and sex) were included for all measures except the social dominance test which included 21 strains (*6 subjects per *Chd8* genotype and sex). Figure adapted from Sittig et al., 2016.

We hypothesized that *Chd8* haploinsufficiency would impact traits across the combined strain population (referred to as *Chd8*^*+/-*^ population effects), and that the severity would differ across genetic backgrounds (referred to as *Chd8*^*+/-*^ strain effects; **Fig. 2)**. Comparison of *Chd8*^*+/-*^ population and strain effects provides estimates of the penetrance of trait disruptions caused by *Chd8* haploinsufficiency across the population and in individual strains. This approach can determine if highly penetrant trait disruptions in human populations with *CHD8* mutations are also highly penetrant in a genetically diverse mouse population, and importantly, identify susceptible and resilient strains. Because each strain is genetically defined and reproducible, identifying the highly penetrant and variable features of *Chd8* haploinsufficiency across and within strains, provides a method to investigate the molecular underpinnings of symptom heterogeneity and discover new treatments. Cohen’s D effect sizes (*d*) reported for *Chd8*^*+/-*^ population and strain effects provide the key representation of significant findings and allow direct comparison of effect magnitudes (large, ≥0.8; medium, ≥0.5; small, ≥0.2).

**Figure 2.**
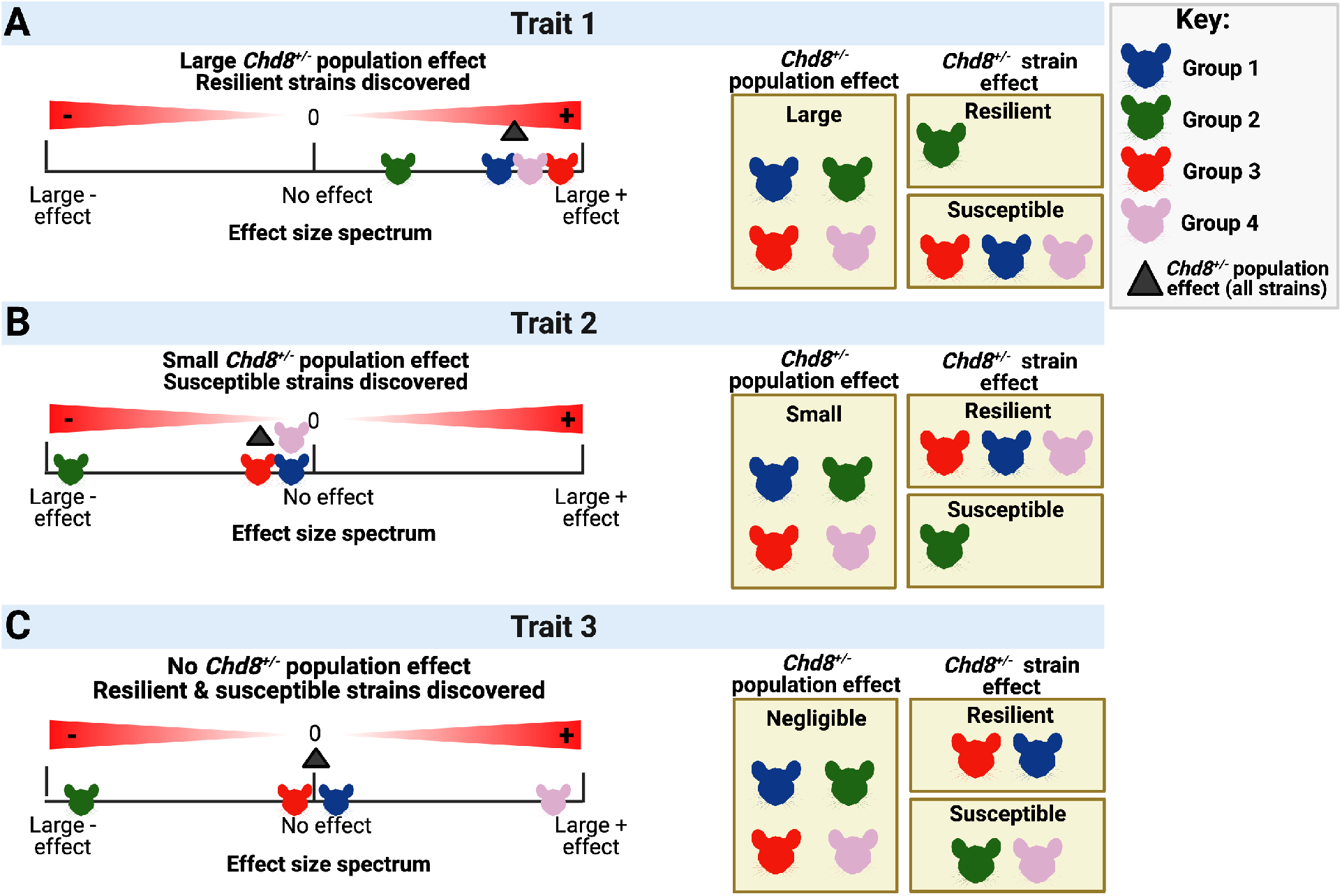
Quantifying population versus individual strain susceptibility to *Chd8* haploinsufficiency. Effect sizes quantify the magnitude and direction of the difference in traits between *Chd8*^*+/-*^ and wild-type (WT) subjects across the genetically diverse strain population (*Chd8*^*+/-*^ population effect, black triangle) and within the same recombinant inbred strain and sex (*Chd8*^*+/-*^ strain effect). Positive effect sizes reflect an increase in *Chd8*^*+/-*^ trait measures compared to WT and negative effect sizes reflect a decrease. *Chd8*^*+/-*^ population effects may differ across traits with large (**A**), small (**B**), or negligible (**C**) effect sizes. Groups of strains may share similarities in how they are impacted across traits (Groups 1-4). If most strains have a large effect size for trait 1, but a few strains have a small effect size, the *Chd8*^*+/-*^ population effect will be large, but the *Chd8*^*+/-*^ strain effect sizes will uncover the resilient strains with the lowest effect sizes. Similarly, if only a few strains show a large strain effect for trait 2, the population effect will be small (B). Further, if there are large *Chd8*^*+/-*^ strain effect sizes but in opposite directions, then the small or absent population effect may mask susceptible strains (C). Strains that are resilient to some traits may be susceptible to other traits.

### Penetrance of trait disruptions caused by Chd8 haploinsufficiency in a genetically diverse population depends on the trait

*Chd8* haploinsufficiency had large to medium population effect sizes for increased social dominance (*d* = 2.33), macrocephaly (*d* = 0.7), and decreased ambulatory behavior in the dark open field (DOF) test (*d* = -0.7) (**Fig. 3A-D**). *Chd8* haploinsufficiency impacted other traits with small to negligible population effect sizes including decreased lifelong body weights, (weaning *d* = -0.4, adolescence *d* = -0.3, young adulthood *d* = -0.3, and terminal *d* = -0.3; **Fig. 3E-H**), increased social sniffing (*d* = 0.4) and aggression (*d* = 0.1) (**Fig. 3G, H**), and increased anxiety-like behavior (*d* = -0.3). Fear learning variables, including fear acquisition, fear expression, and fear extinction were not significantly different between the *Chd8*^*+/-*^ and WT populations and generated negligible effect sizes (*d* = 0.1; **Fig. 3L-N**).

**Figure 3.**
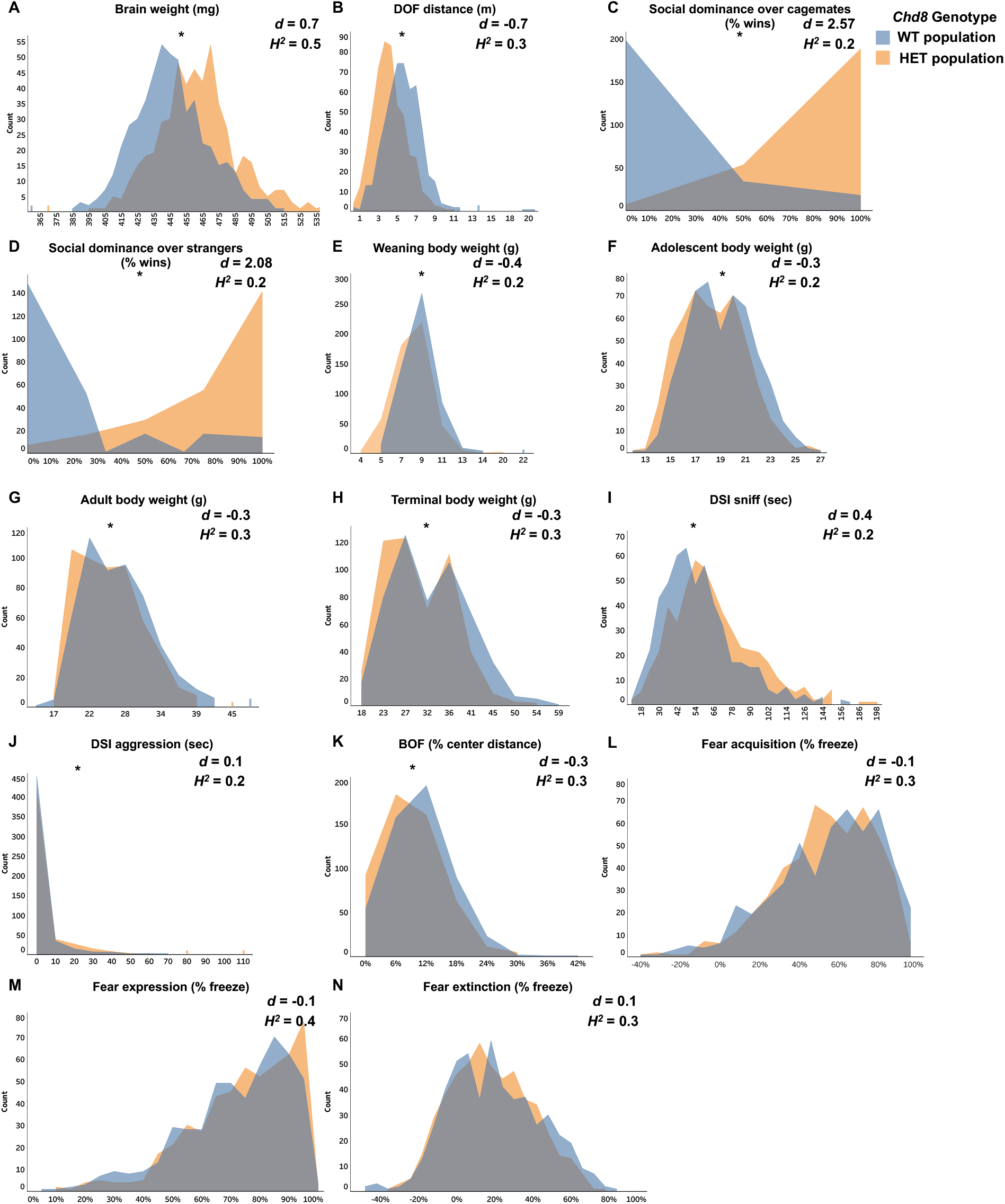
Population-based penetrance of phenotypic differences caused by *Chd8* haploinsufficiency differs across traits. The median population distribution of traits shifted as a function of genotype for wild-type (WT; blue) and *Chd8*^*+/-*^ (HET; orange) males and females from 33 strains (n = 1,051). Gray reflects overlapping WT and HET population distributions shown for brain weight (**A**; U= 82,140, p < 0.001), distance traveled in the dark open field (DOF) test (**B**; U= 191,343, p < 0.001), social dominance over opposite genotype cagemates (**C;** 21 strains: U=4399, p < .001) and strangers (**D;** 21 strains: U= 5694, p < .001), weanling body weight (**E**; U= 169,334, p < 0.001), adolescent body weight (**F**; U= 156,371, p < 0.001), adult body weight (**G**; U= 153,917, p < 0.001), terminal body weight (**H**; U= 152,971, p < 0.001), duration of sniffing a same-sex WT B6 juvenile in the direct social interaction (DSI) test (**I**; U= 105,271, p < 0.001), aggression duration in the DSI test (**J**; U= 124514, p < 0.05), and percent center distance traveled in the bright open field (BOF) test (**K**; U= 158549, p <0.001). Percent time freezing during fear acquisition (**L**; U= 130309, p = 0.18), fear expression (**M**; U = 126,341, p= 0.11), and fear extinction (**N**; U= 138,703, p = 0.345) were not different between *Chd8* genotype populations. Asterisks indicate significant (*i*.*e*., p < 0.05) median shifts between *Chd8* genotype groups from Mann-Whitney U tests. Cohen’s D effect sizes (*d*) and heritability estimates (*H*^*2*^) are indicated for each trait. Negative effect sizes indicate a decrease in the *Chd8*^*+/-*^ population trait mean compared to WT.

There were also population differences between sexes in traits as well as the impact of *Chd8* haploinsufficiency (**SFig. 1**). Female brains weighed more than males, and whereas *Chd8*^*+/-*^ males and females had significantly larger brains than WTs, the mutation effect size was greater in males (d = 0.9) compared to females (d = 0.5). Moreover, compared to females, males had larger body weights, spent more time sniffing a same-sex B6 conspecific in the DSI test, were less anxious in the BOF test, and had higher fear extinction scores. In contrast, females had higher fear acquisition and fear expression scores. Lastly, WT females traveled farther than WT males in the DOF test, and *Chd8*^*+/-*^ males had higher aggression in the DSI task than *Chd8*^*+/-*^ females.

### *Chd8* haploinsufficiency impacts variation between traits

To capture the shared population variance across many traits, they were reduced into principal components (PCs) by performing a Principal Components Analysis (PCA). The impact of *Chd8* haploinsufficiency on PCs in the population and by strain and sex was then assessed (**Fig. 4A)**. Social dominance scores were not included in the PCA because not all strains were tested (N=12 strains not tested). The 5 extracted PCs revealed expected relationships between traits in the population including body weights (PC1), social behavior measures (PC4), and fear learning variables (PC5). Moreover, DOF ambulatory behavior and BOF anxiety-like behavior were negatively associated with fear expression (PC2). In addition, PC3 revealed relationships between weaning body weights, terminal brain weights and BOF anxiety-like behavior.

**Figure 4.**
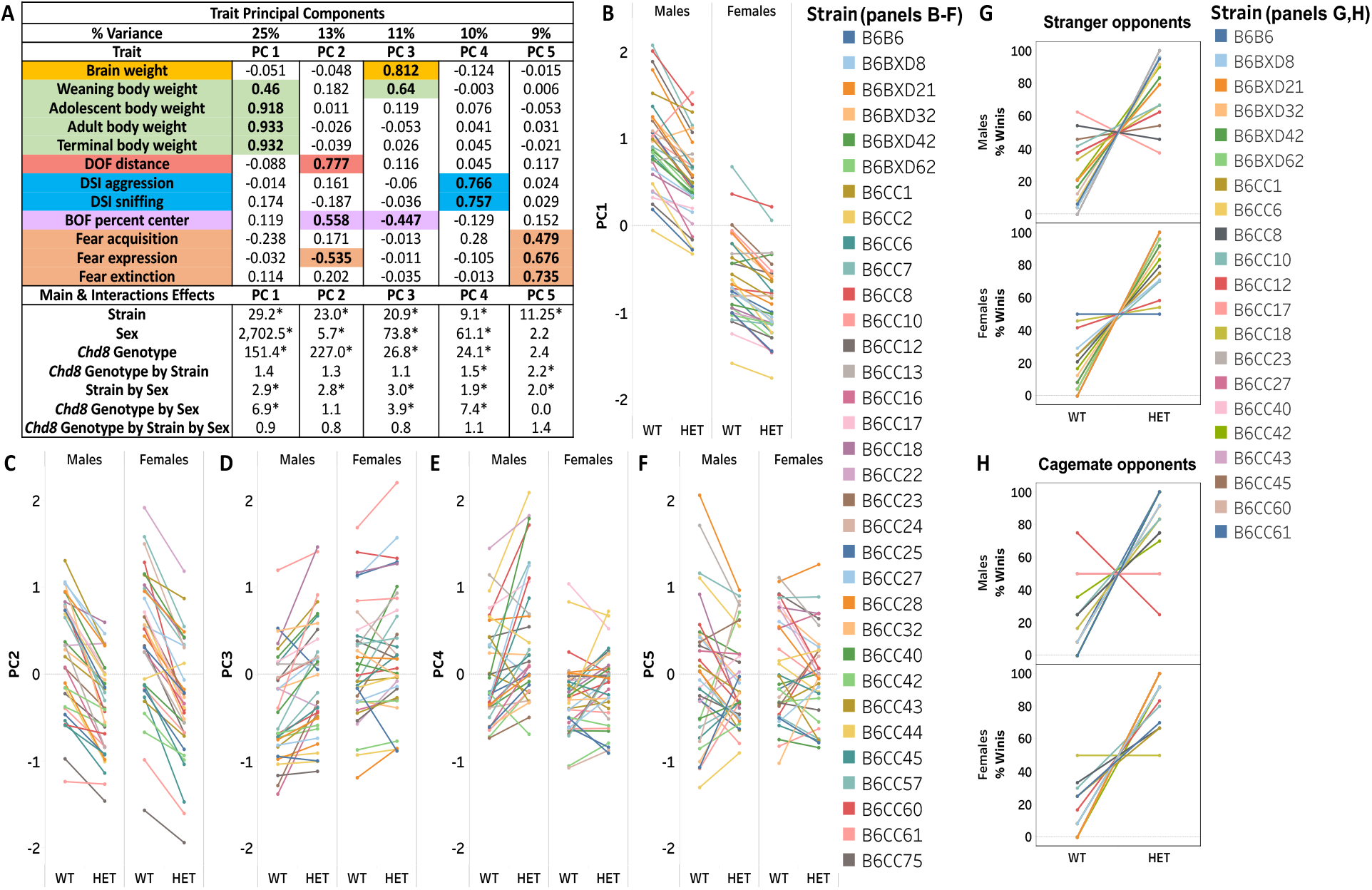
Trait relations and the impact of *Chd8* haploinsufficiency on trait principal components and social dominance. A rotated component matrix with Varimax rotation and Kaiser normalization extracted 5 principal components (PCs) that capture 68% of the trait variance (**A**). The percent variance for each PC with loading coefficients are listed in the table (A). Bold numbers indicate coefficients greater than 0.4, reflecting traits that highly covary within a PC. Traits are color-coded by similarity to each other along with their respective high loading coefficients. The F statistic from 3-way ANOVA for each PC is reported and significance (*i*.*e*., p < 0.05) is indicated by asterisks (A). Data points in the line graphs **B-H** represent the average PC score (B-F) or percent social dominance wins (G, H) by sex, strain and *Chd8* genotype. Lines connect *Chd8* wild-type (WT) and *Chd8*^*+/-*^ (HET) males and females per strain and colors represent different strains. A larger PC score in one strain compared to another indicates larger trait values in that strain while the slope and direction of the lines connecting the WT and *Chd8*^*+/-*^ indicate the magnitude and direction of the effect for each strain. 33 strains were included in the PCA (A-F) and 21 strains were tested for social dominance (G, H; 3-way *Chd8* genotype by strain by sex interaction G, F_1, 498_ = 4.364, P < 0.001; H, F_1, 498_ = 6.487, P < 0.001).

Decreased PC1 scores in the *Chd8*^*+/-*^ population indicated reduced body weights compared to WT, with the effect stronger in males than females. Reduced PC2 scores in the *Chd8*^*+/-*^ population also revealed reduced ambulatory behavior and increased anxiety-like and fear expression (**Fig. 4B, C**), which at the strain level may reveal some for which there is greater anxiety-like and fear behaviors and for others driven by differences in activity. Moreover, increased PC3 scores in *Chd8*^*+/-*^ mice indicated increased anxiety-like behavior and brain weights compared to WTs, which covaried only with weights at weaning but not at other times, and the effect was stronger in males than females **(Fig. 4D)**. Increased PC4 scores in *Chd8*^*+/-*^ males, but not females, reflected increased DSI aggression and sniffing compared to WTs **(Fig. 4E)**. There was no *Chd8*^*+/-*^ population effect on PC5 scores, but *Chd8*^*+/-*^ mice from some strains had decreased scores whereas *Chd8*^*+/-*^ mice in other strains had increased scores compared to WTs, reflecting bidirectional manifestations of *Chd8* haploinsufficiency on fear learning across strains (**Fig. 4A, F**). There was striking variation amongst strains in score differences between WT and *Chd8*^*+/-*^ for all PCs but this was particularly heterogeneous for PCs 3-5. Similarly, while *Chd8*^*+/-*^ males and females across strains won more social dominance matches than WTs **(4G, H**), there was substantial variation between individual strains and sexes, including in the occurrence, magnitude, and direction of differences.

### Strain and sex modify penetrance of *Chd8* haploinsufficiency across every trait

As a correlate to assessing symptom severity across a clinical population, for each trait we examined the effect size distribution for individual strains. Remarkably, every trait, including those with large to negligible population effect sizes, exhibited a large range of *Chd8*^*+/-*^ strain effects including large and negligible strain effect sizes (**Fig. 5A**).

**Fig 5.**
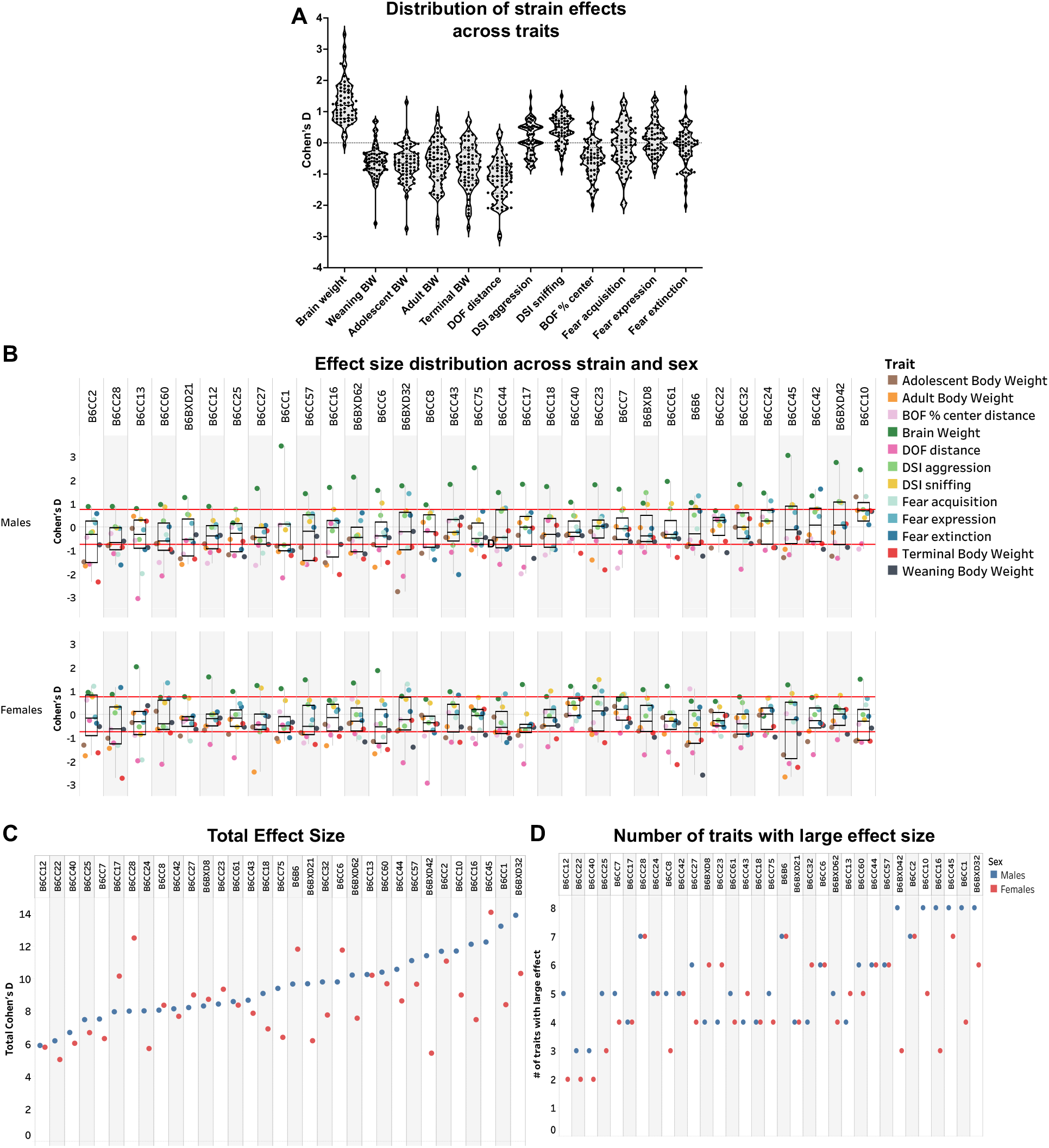
Severity of *Chd8* haploinsufficiency trait disruption varies broadly for every trait and is modified by strain and sex. The distribution of *Chd8*^*+/-*^ strain effect sizes varies across traits (N = 66 strain and sex groups per trait; **A**). *Chd8*^*+/-*^ strain effect size distributions across 12 traits within individual strains and sexes revealed marked heterogeneity in the combination of specific traits that were impacted (**B**). Traits are represented by colors. Strains are ordered in ascending trait means for males. Red lines highlight effect sizes below and above a large effect (*i*.*e*., |*d*| = 0.8). The sum absolute value of effect sizes across 12 traits for males (blue) and females (red) varies between strains and sexes (**C**, plotted in ascending order for males). The number of traits on which strains and sexes are impacted with large effect sizes also varies (**D**). Strains in panel D are graphed in the same order as panel C to facilitate comparison between graphs. BW = body weight.

As a correlate to assessing symptom severity within an individual, for each strain we examined the effect size distribution across traits. For most strains, there was marked heterogeneity on trait impact, with the specific strain and sex that had the lowest and highest effect sizes differing across all traits (**Fig. 5B**). The combination of specific traits that were impacted consistently differed between strains and sexes, underscoring the complexity of trait alteration and highlighting how genetic modifiers might interact with *Chd8* heterozygosity to influence clinical heterogeneity.

To better understand differences in how the strains and sexes were impacted across many complex traits, absolute values of Cohen’s D effect sizes for every trait, except social dominance, were summed into a total effect size score per strain and sex **(Fig. 5C)**. Total effect sizes ranged from a minimum of 5 in B6-CC22 females to a maximum of 14 in B6-BXD32 males and B6-CC45 females **(Fig. 5C)**. Sex as a driver was noteworthy, with sex differences in total effect size for many strains **(Fig. 5C)**. Next, to identify how many traits, strains and sexes were impacted with a large effect size, the number of traits with a large Cohen’s D (i.e., |d| > 0.08) were summed across 12 traits (**Fig. 5D)**. It was most common for males and females in approximately 27% of strains to have large effect sizes across 5 and 4 traits (33-42%), respectively, although the specific traits and strains that were impacted differed between sexes (**Fig. 5D**). The least impacted strains had large effect sizes across 3 traits in males (6% of strains) and 2 traits in females (9% of strains). The most impacted strains had large effect sizes across 8 traits in males (18% of strains) and 7 traits in females (12% of strains) (**Fig. 5D)**.

B6-CC22 males and females had some of the lowest effect sizes with a large effect size across only 2 traits in females and 3 traits in males **(Fig. 5C, D)**. B6-CC45 males and females were impacted with a large effect size across 8 traits in males and 7 in females. However, the combination of traits on which males and females were impacted differed.

### *Chd8* haploinsufficiency induced co-occurring trait disruptions across the population and amongst strains and sexes

To first identify if there were patterns in the combination of traits impacted by *Chd8* haploinsufficiency in the population, an exploratory factor analysis (EFA) on *Chd8*^*+/-*^ strain effect sizes across 12 traits in males and females was performed. In addition, hierarchical clustering analysis (HCA) was performed, and Spearman’s correlation coefficients extracted (**Fig 6)**. Results indicated consistent impact of *Chd8* haploinsufficiency across body weight trajectories (FS1). In addition, the impact of *Chd8* haploinsufficiency on DSI aggression and sniffing coincided with fear acquisition and expression (FS2). Weaning body weight effect sizes covaried both negatively (FS3) and positively (FS4) with BOF effect sizes, indicating two distinct relationships between the impact of *Chd8* haploinsufficiency on weaning body weights and anxiety-like behavior in the population. Fear expression effect sizes were correlated with fear acquisition and extinction but loaded independently on FS5 indicating that the impact of *Chd8*^*+/-*^ on fear expression is independent of the impact of *Chd8* haploinsufficiency on other traits. *Chd8*^*+/-*^ strain effects for brain weight and DOF distance did not significantly correlate nor covary with other traits reflecting the high penetrance of *Chd8* haploinsufficiency on macrocephaly and activity across strain and sex.

**Figure 6.**
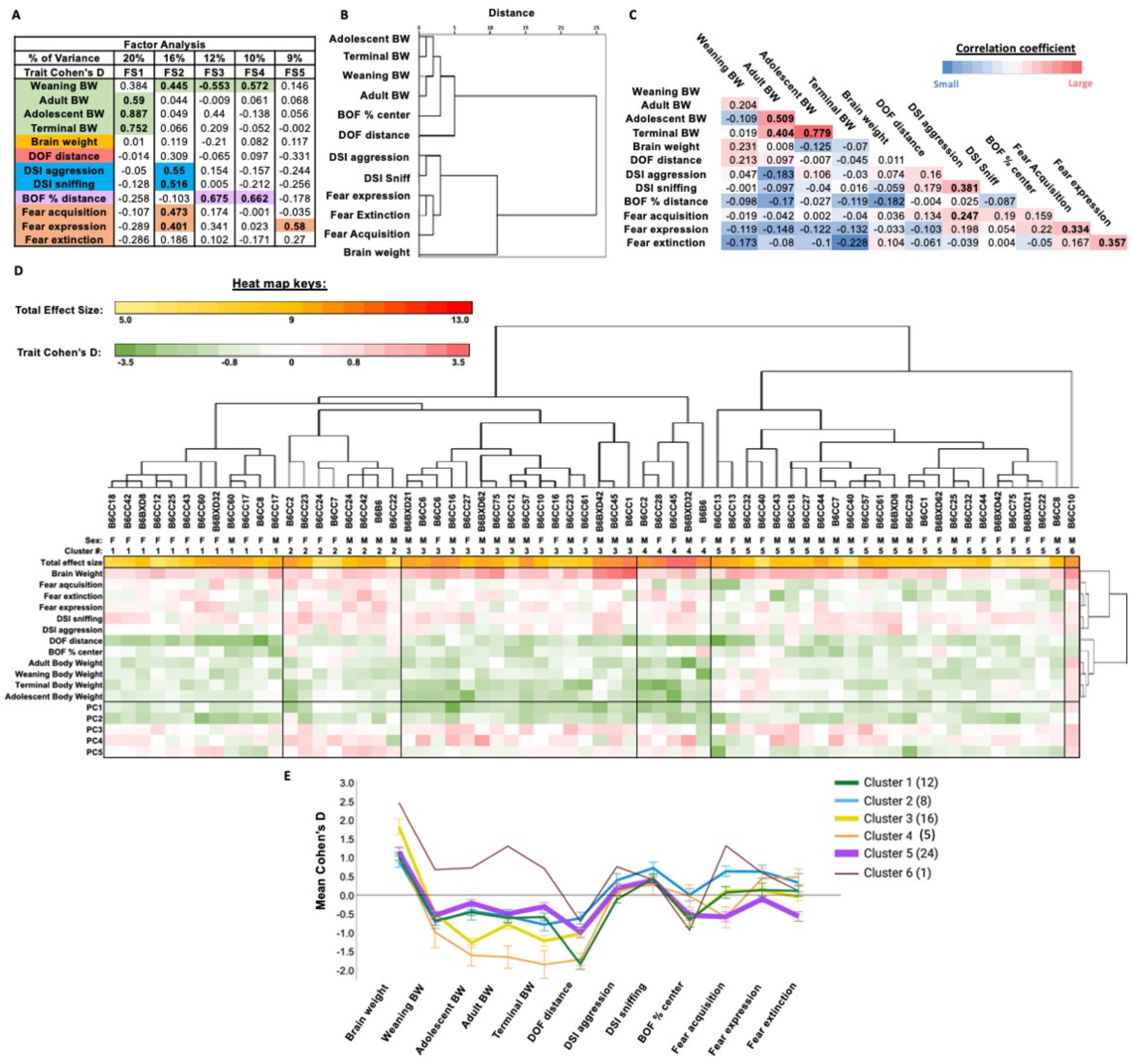
Genetic background regulates the covariance of traits impacted by *Chd8* haploinsufficiency across a population. Exploratory factor analysis (EFA) on *Chd8*^*+/-*^ strain effect sizes resulted in five factor scores (FS; **A**). High loadings *(i*.*e*., *>* 0.4) are bolded and color-coded with the corresponding trait. Hierarchical cluster analysis (HCA) on *Chd8*^*+/-*^strain effects produced congruent results to EFA (**B**). Significant (*i*.*e*., p < 0.05) Spearman’s correlation coefficients for *Chd8*^*+/-*^ strain effects across traits are bolded (**C**). Heatmaps of *Chd8*^*+/-*^ strain effects summed across 12 traits (total effect size) as well as across individual traits and principal components (PC 1-5) with strains and sexes ordered by their similarities as determined by HCA (**D**). The dendrograms represent the relation between *Chd8*^*+/-*^ strain effects across traits (Y axis) and strain and sex groups (X axis). Effect size means (+/- SEM) for each cluster varies across traits (**E**). Different line colors represent different cluster groups. Line thickness and numbers in the legend reflect the number of strain and sex groups in each cluster. FS = factor score; BW = body weight; DOF = dark open field; DSI = direct social interaction task; BOF = bright open field

To classify strains and sexes by similarities in effect sizes across traits, an HCA was performed on *Chd8*^*+/-*^ strain effects in males and females. Next, heatmaps of *Chd8*^*+/-*^ strain effects across traits and PCs were constructed to visualize the impact of *Chd8* haploinsufficiency across traits within and between cluster groups (**Fig. 6D)**. The dendrogram identified 6 main clusters, each differing in size, sex composition, and average effect sizes. Within clusters, strains and sexes shared effect sizes for some traits whilst there was heterogeneity amongst other traits.

Cluster 5 (54% males) was the largest (36% of strains and sexes) as well as the most resilient cluster, with the lowest cluster effect size mean. Cluster 5 was characterized most notably by large effect sizes for macrocephaly and decreased DOF distance in *Chd8*^*+/-*^ compared to WT mice. *Chd8*^*+/-*^ mice in cluster 5 also had decreased weaning and adult body weights, fear acquisition and extinction, and increased anxiety-like behavior with medium effect sizes on average (**Fig. 6E**).

Cluster 3, the second largest cluster (24% of strains and sexes, 69% male), differed from cluster 5 by being particularly susceptible (i.e., having very large effect sizes) to macrocephaly and decreased body weights after weaning in *Chd8*^*+/ -*^ mice compared to WT.

Cluster 1 was the third largest cluster (18% of strains and sexes, 2% males) that were amongst the most susceptible for decreased DOF ambulatory behavior in *Chd8*^*+/ -*^ mice. Clusters 1 and 3 had negligible effect sizes for fear learning variables.

Cluster 2 consisted of 12% of strains and sexes (50% male) that were amongst the most impacted for increased DSI sniffing and the least impacted for decreased DOF ambulatory behavior in *Chd8*^*+/ -*^ mice. In addition, cluster 2 had negligible effect sizes, on average, for BOF anxiety-like behavior and medium effect sizes for *increased* fear learning in *Chd8*^*+/ -*^ mice.

Cluster 4 comprised 8% of strains and sexes (40% male) with the largest cluster effect size average that were severely impacted on body weights and ambulatory behavior but unimpacted on BOF anxiety-like behavior.

Remarkably, cluster 6 consisted only of B6-CC10 males, exhibiting the largest cluster effect size for macrocephaly. B6-CC10 males were the only strain and sex group to have *increased* body weights in *Chd8*^*+/-*^ compared to WT males and with large effect sizes. B6-CC10 *Chd8*^*+/-*^ males also had increased BOF anxiety-like behavior compared to WT as well as increased DSI aggression, with large effect sizes.

## Conclusions

A genetically wholistic understanding of human disease is essential for classifying the symptom profiles that are highly penetrant and identifying the genetic backgrounds that modify their presence and severity. This is particularly true for highly heritable but genetically heterogenous disorders like ASD, for which recent studies highlight the role of both common and rare genetic variants, and their interaction, in disorder risk and severity (1,2). Analyses of *CHD8* patient-derived iPSC neurons highlight the important role for donor-genetic background in determining how *CHD8* haploinsufficiency affects the development of inhibitory and excitatory neural lineages, which is relevant for understanding individual differences in cortical excitation and how this correlates with clinical symptom severity (25,26). Complementing such results, our data demonstrate that the systematic introduction of genetic diversity (common and rare) into a B6/inbred mouse model of *Chd8* haploinsufficiency can uncover the range of trait disruptions in a genetically diverse population in addition to susceptible and resilient individual and groups of strains. The core phenotypes of human *CHD8* haploinsufficiency were recapitulated in the mouse population, including high prevalence of macrocephaly and social behavioral changes with higher variance in occurrence and severity of co-occurring traits like anxiety and learning deficits. Sex is an important modifier of trait penetrance, as is often observed in ASD, but often unknown for rare mutations. We also found evidence that traits covary themselves in addition to their disruption by *Chd8* haploinsufficiency and classified groups of strains and sexes with shared patterns of trait disruption occurrence, magnitude, and direction. This provides for the first time a curated list of CC and BXD strains and sexes that are openly available and can serve as genetic and molecular anchor points for the study of other high confidence NDD genes, and mechanistic discoveries on the origins of susceptibility, resilience, trait covariance, and responses to treatments. From this foundation comes the promise of discovering NDD etiologies and treatment discovery at both population and individual levels of analyses.

## Acknowledgments

The authors thank Amanda Whipple and Patricia Aguiar for their assistance with behavioral testing, genotyping, breeding, and colony management. We also thank Sunny Lee, Shreya Harappanahally, Shrestha Vijayendra, and Khalifa Elmagarmid for their assistance in scoring behavioral videos. We are grateful to Rob Williams, Kathie Eagleson, Brian Dias, and Rex Moates for their helpful feedback on versions of this manuscript.

## Funding

National Institute of Mental Health grant R21MH118685 (P.L., A.K.)

National Science Foundation Postdoctoral Research Fellowship in Biology grant 2011039 (M.T.), The Saban Research Institute Research Career Development Fellowship (M.T.), Simms Mann Chair in Developmental Neurogenetics and Developmental Neuroscience and Neurogenetics Program, Children’s Hospital Los Angeles (P.L.), and WM Keck Chair in Neurogenetics, Keck School of Medicine of the University of Southern California (P.L.).

## Author contributions

Conceptualization: PL, AK, MT

Methodology: PL, AK, MT

Investigation: AK, MT, PL

Visualization: MT, AK, PL

Funding acquisition: PL, AK, MT

Project administration: AK, MT, PL

Supervision: PL, AK, MT

Writing – original draft: MT, PL, AK

Writing – review & editing: MT, PL, AK

## Competing interests

Authors declare that they have no competing interests.

## Data and materials availability

Datasets used in the analysis are available on GeneNetwork.org

## Supplementary Materials

Supplementary Text

Fig. S1

Data S1 to S2

## Notes

### Competing Interest Statement

The authors have declared no competing interest.

