## Supplemental for "Mouse population genetics phenocopies heterogeneity of human *Chd8* haploinsufficiency"

Materials and Methods

All experimental procedures were approved by the University of Southern California (USC) Institutional Animal Care and Use Committee under protocol 11844-CR011. In addition, all experimental procedures followed the Guidelines for the Care and Use of Laboratory Animals by the National Institutes of Health.

Mice

Mice were housed in the Ray R. Irani vivarium at the USC main campus from 2019-2021. Mice were housed in standard ventilated cages on a 12 h light/dark cycle (lights on at 6:00 AM) in a temperature (20–22°C) and humidity (40-60%) controlled room with *ad libitum* access to standard rodent chow and filtered water. C57Bl/6J (B6) mice that were heterozygous for *Chd8* (*Chd8^+/-^* ) were originally received from the laboratory of Dr. Feng Zhang (29) and maintained in our colony at USC. Males from 27 Collaborative Cross (CC) and 5 BXD recombinant inbred strains, in addition to C57BL/6J (B6), were obtained from The Jackson Laboratory (Bar Harbor, ME) at 6 to 8 weeks of age and allowed to habituate to the colony for two weeks prior to paring with B6-*Chd8^+/-^* females. CC and BXD strains were chosen based off health data provided for strains by the Collaborative Cross and Complex Traits Consortium in addition to prior work from our lab demonstrating variability in sociality in a BXD panel. To control for differences in paternal care, sires were removed prior to litter birth. The experimental F1 WT and *Chd8^+/-^* B6-CC, B6-BXD, and B6-B6 male and female littermates were weaned at P21 and housed with same sex cagemates. At weaning, tail snips (~1-2mm) were collected for *Chd8* genotyping. Experimental mice were tested in three cohorts over 2 years. The first cohort included 17 strains, the second cohort included 12 strains and the last cohort included 4 strains. Power calculations based on our previous studies indicate 80% power to detect an effect size of 40% with an N of 6 per *Chd8* genotype and sex group. To increase the power and sensitivity, we included 8 subjects per *Chd8* genotype, sex, and strain group in this study. Subjects were tested in the following order with at least a 1 week break between tests: DOF, DSI, BOF, and cued fear conditioning. The subset of 21 strains that were tested in the social dominance tube test were tested the week prior to fear conditioning.

Behavioral tests were conducted during the light cycle (between 6:00 AM and 5:00 PM). Mice were transported to the testing room, or a holding room adjacent to the testing room, at least 45 minutes prior to testing. Body weights were taken at weaning (P21), adolescence (P35), as young adults directly after the DSI test (~P120), and as older adults directly prior to euthanasia (~P192). A total of 1,051 mice entered the study. Researchers were blind to the genotype of the subjects during behavioral testing and body and brain weighing. The order of strains going through behavioral testing was randomized. Males and females were tested in separate batches so that both sexes were not occupying the same behavioral suite during any test and batches were separated by about a day. Once entered in the study subjects were included for the full duration of the study, but some subjects were excluded due to unexpected expiration or pronounced defects like malocclusion. There were not enough exclusions to statistically analyze strain or genotype effects, but there did not appear to be exclusions specific to any one strain.

Dark Open Field (DOF) Test

Baseline activity levels were assessed using the DOF test. In this task, mice were placed into a dark Plexiglas testing arena measuring 27.31 L x 27.31 W x 20.32 H cm (Med Associates, Inc.) and that was enclosed in a larger cabinet to ensure darkness during the 30-minute test. Activity levels of the freely moving subjects were captured with laser sensors fixed throughout the box that were fed into a computer with Med Associates activity tracker software installed. Data for each run was later extracted from the activity tracker software for further analysis. Distance traveled was the main dependent variable analyzed for the DOF test in this study.

Direct Social Interaction (DSI) Test

Social behaviors towards a same-sex conspecific were assessed in the DSI test as described previously (15). Subjects were placed in the rectangular Plexiglass testing chamber (30 L X 19” W X 19 H cm) for a 10-minute habituation period. Then, a B6 juvenile (P26-P30; mean P28) of the same sex as the subject was placed into the testing chamber for the 6-minute test. Juvenile B6 males and females were used in the DSI test to minimize potential aggressive behaviors. Behavior was videotaped from top-down and frontal viewpoints. Behavioral videos were later scored by trained researchers blind to subject genotype with Boris, an open-source behavioral scoring software (30). Behaviors scored included durations and frequencies for sniffing, aggression, auto-grooming, mounting, no behavior, and tail rattling. Sniffing was defined as the subject’s nose being approximately 1 cm away from the juvenile and sniffing anywhere on the juvenile’s body, including their tail. Aggression was scored when it became overt and included biting, dragging, tumbling, and forceful pushing. No behavior was the default behavior scored when subjects were not engaged in other defined behaviors. Mounting was scored when the subjects were on their hindlegs with their forepaws extended and a hunched posture over any part of the juvenile’s body. Only 1.4% of mice across 10 (3%) strains displayed mounting. Therefore, mounting was excluded from further analyses.

Bright Open Field (BOF) Test

Anxiety-like behaviors were assessed with the BOF test. The BOF test began when subjects were placed into the center of a brightly illuminated Plexiglas test chamber measuring 27.31 L x 27.31 W x 20.32 H cm (Med Associates, Inc.). The test chamber was enclosed in a larger cabinet during the 30-minute test to ensure an isolated environment and reduced noise. Location of the mouse was tracked by series of laser sensors fixed throughout the box and transmitted to Med Associates activity tracker software. The tracking accuracy of the sensors was verified by more than one researcher repeatedly throughout this study. The distance traveled the outside 2” perimeter and the distance traveled in the center of the box were later extracted from the activity tracker software for further analysis of the percent center distance traveled.

Social Dominance Tube (SD) Test

*Chd8^+/-^* and WT mice from a subset of strains (N=21 strains tested) of the same sex and strain were paired in the SD test. The SD test began when a *Chd8^+/-^* and WT mouse simultaneously entered the opposite ends of a narrow tube. Two researchers that were blind to *Chd8* genotype coordinated the removal of each mouse from the home cage and placement into the opposite openings of the tube with their noses oriented inside the tube openings until the mice entered and met in the approximate middle of the tube. The mouse that was the first to leave the tube was recorded as the “loser” and the mouse remaining in the tube was recorded as the “winner”. “Winners” that did not leave the tube following the “losers” exit were coaxed to continue through out of the tube by gentle nudging of their backside with a flexible rubber rod. Prior to SD testing, mice were trained to run through the tube approximately 10 times for two consecutive training days. Mice were trained to run through the tube by consistently placing their nose into the tube opening until they advanced into the tube and then gently nudging their backside with a rubber rod to coax them to continue through the tube. Once mice entered the tube, they were generally willing to continue entering freely without researcher interference, but they often preferred to remain in the tube and therefore were nudged through the tube. Tube sizes were either small, medium, or large and were picked to best fit each strain so that a mouse could not turn around to exit the tube, exiting only by moving forward or backing out of the tube. All SD matches reported in this study were between *Chd8^+/-^* and WT same sex and strain conspecific. Males and females were first tested against unfamiliar conspecifics from different cages and then against their opposite *Chd8* genotype cage mates. Each subject went through 4 matches between opposite *Chd8* genotype strangers and 2 matches between cage mates.

Fear conditioning

To assess learning and memory, cued fear conditioning was conducted in an automated near-infrared video fear conditioning system (Med Associates) as described previously (16). The testing chamber (30 L X 25 W X 21 H cm) had stainless steel walls and floor bars and a transparent acrylic door. The fear conditioning test encompassed 4 days and included habituation on day 1, training on day 2, memory testing on day 3, and memory extinction (*i.e,* memory persistence) testing on day 4. On habituation day 1, mice were acclimated for 30 minutes to the test chamber. On training day 2, 5 presentations of a 5 kHz, 85dB, 30-second tone (conditioned stimulus; CS) were paired with a 0.5 mA, 2 second foot-shock unconditioned stimulus (US). The CS and US co-terminated. The first CS-US presentation took place 180 seconds after the start of the test in addition to between each CS-US presentation. On test day 3, cued-fear was measured approximately 24 hours later in a novel context. Textured clear plastic walls and white smooth plastic floor inserts provided a novel context, and subjects were presented with 10 CS presentations 30 seconds long with 60 seconds in between CS intervals. Approximately 24 hours later, on the last testing day, the persistence of fear memory was tested in the same manner as on testing day 3. Fear conditioning tests were videotaped (30 frames/s) under near-infrared light and freezing times were scored automatically using VideoFreeze software (Med Associates). Freezing times were defined as no movement for a duration of 1 second (30 frames). Testing chambers were cleaned by first spot cleaning waste with Kim wipes followed by wiping down with paper towels with 70% ethanol. Lastly, all testing materials were wiped down with filtered water and then dried thoroughly.

Learning during training (fear acquisition) was calculated by subtracting the percent time the subject spent freezing for the 28 seconds directly prior to the first shock presentation (freezing to the tone alone defined as baseline percent freezing) from the percent time freezing during the last (5^th^) presentation of the CS on training day. Learning during testing (fear expression) was calculated by averaging the percent time the subject spent freezing during the first three presentations of the CS on testing day. Fear extinction scores were calculated by subtracting the mean percent freezing to the CS during testing day from the average percent time freezing to the last three presentations of the CS on the last testing day.

Body and brain weights

Body weights were taken of all subjects at weaning (P21) and at adolescence (P35). Adult body weights were taken directly after the DSI test (mean = P125 +/- 14) and at euthanasia directly prior to brain removal (mean = P 192 +/- 13).

Mice were euthanized with vaporized (~4%) isoflurane exposure, and, upon cessation of breathing, death was confirmed by decapitation. Directly after isoflurane exposure but prior to decapitation, subjects body weights were recorded, and confirmatory tail snips (~4 mm) were collected to confirm *Chd8* genotypes. Brains were extracted immediately and brain weights obtained. Two researchers, experts at dissecting brains, conducted all brain dissections, and the entire process from the point of euthanasia to weighing the brain took approximately 10 minutes per subject.

Statistics

Traits analyzed include weaning, adolescent, young adult, and terminal body weights, brain weight, DOF distance traveled, DSI aggression and sniffing durations, percent distance traveled in the center of the BOF, fear acquisition, fear expression, and fear extinction for all 33 strains. The percentage of wins during the SD test between opposite *Chd8* genotype cagemates and strangers was also analyzed for 21 strains.

The impact of *Chd8* heterozygosity on trait distributions across strains was investigated with parametric and non-parametric tests including R^2^ values, Mann-Whitney U statistics and probability values in addition to group median differences with 95% confidence intervals. In addition, parametric and non-parametric effect size estimates were calculated including Cohen’s D with 95% confidence intervals, Vargha & Delaney’s A probability of stochastic superiority (31), and the probability or common language effect size (32). Statistics for the strain population, with sexes combined and separated, are listed in SData1 and by strains and sex groups in SData2. All statistical analyses and calculations were conducted in SPSS and Excel. Figures and graphs were constructed with Tableau, Prism, and Bio Renderer.

Supplementary Text

Spread of data: Outliers, normality, and homogeneity of variance

Histograms of traits by genotype indicated no outliers in the WT or *Chd8^+/-^* populations that were not indicative of strain differences. Considering a major goal of this study is to identify strains with extreme phenotypes (i.e., resilient or susceptible), outliers were further assessed per strain rather than the entire population. Z-scores across traits by strain and trait were all below 3.

Normality was evaluated by *Chd8* genotype group with sexes combined and separated. Skewness and Kurtosis values as well as Kolmogorov-Smirnov and Shapiro-Wilk test statistics and significance (SData1), in addition to trait histograms (Fig. 3), revealed both approximately normal and non-normal distributions across traits.

Homogeneity of variance across traits, strains, and sexes was analyzed with coefficients of variation (CoV) and Levene’s test (SData1). CoV was calculated by dividing the strain, sex, and genotype group standard deviation by the group mean. In the cases in which there were no instances of a trait displayed in a group, a CoV of 0 was assigned for all groups to be included for comparison (i.e., if no subjects in a group displayed aggression during the DSI test, then a CoV of 0 was assigned to that group). Variation in CoV values was not equal between *Chd8* genotype groups across traits; *Chd8^+/-^* had lower overall CoV scores across traits compared to WT (WT CoV mean = 0.427; Chd8 Het = 0.358). However, when sex was considered, HOV was equal between *Chd8* genotype groups for most of the traits except for brain weight, DSI aggression duration, fear acquisition, and fear extinction in males in addition to weaning body weight in females (SData1). Some traits were associated with lower CoV scores and less variability between strains compared to other traits that had large CoV scores and large differences between strains within a trait. Strains had lower CoV scores for body and brain weights, in addition to less variation between strains, while behavioral variables like DSI aggression and fear extinction consisted of higher CoV scores across strains with more variation between strains.

Heritability

Broad-sense heritability (H^2^) values were calculated as described previously (16), using one way-ANOVA to determine the proportion of phenotypic variance accounted for by strain in the WT population. The most heritable traits were brain weight and terminal body weight in males and females (H^2^ = 0.50 and 0.6 in males and 0.65 and 0.71 in females, respectively). Adolescent and adult body weights were also highly heritable in both sexes followed by DOF distance, fear expression and fear extinction. H^2^ scores by genotype with sexes combined and separated are listed in SData 1.

Principal Component Analysis

A rotated component matrix on z-scores across 12 traits for each subject with Varimax rotation and Kaiser normalization converged in 6 iterations and extracted 5 principal components (PCs; **A**). Kaiser-Meyer-Olkin Measure of Sampling Adequacy was 0.711 and Bartlett’s Test of Sphericity was X^2^ (55, 1,041) = 3417.701 (p < 0.001).

Factor Analysis and Hierarchical Clustering Analysis

Exploratory factor analysis (EFA) on Cohen’s D effect sizes for 12 traits across strain and sex groups (N=66) with the principal axis factoring extraction method resulted in five factor scores (FS; **A**). Hierarchical cluster analysis (HCA) on trait Cohen’s D values with agglomeration schedule, proximity matrix, Ward’s linkage and squared Euclidian distance produced congruent results to EFA.

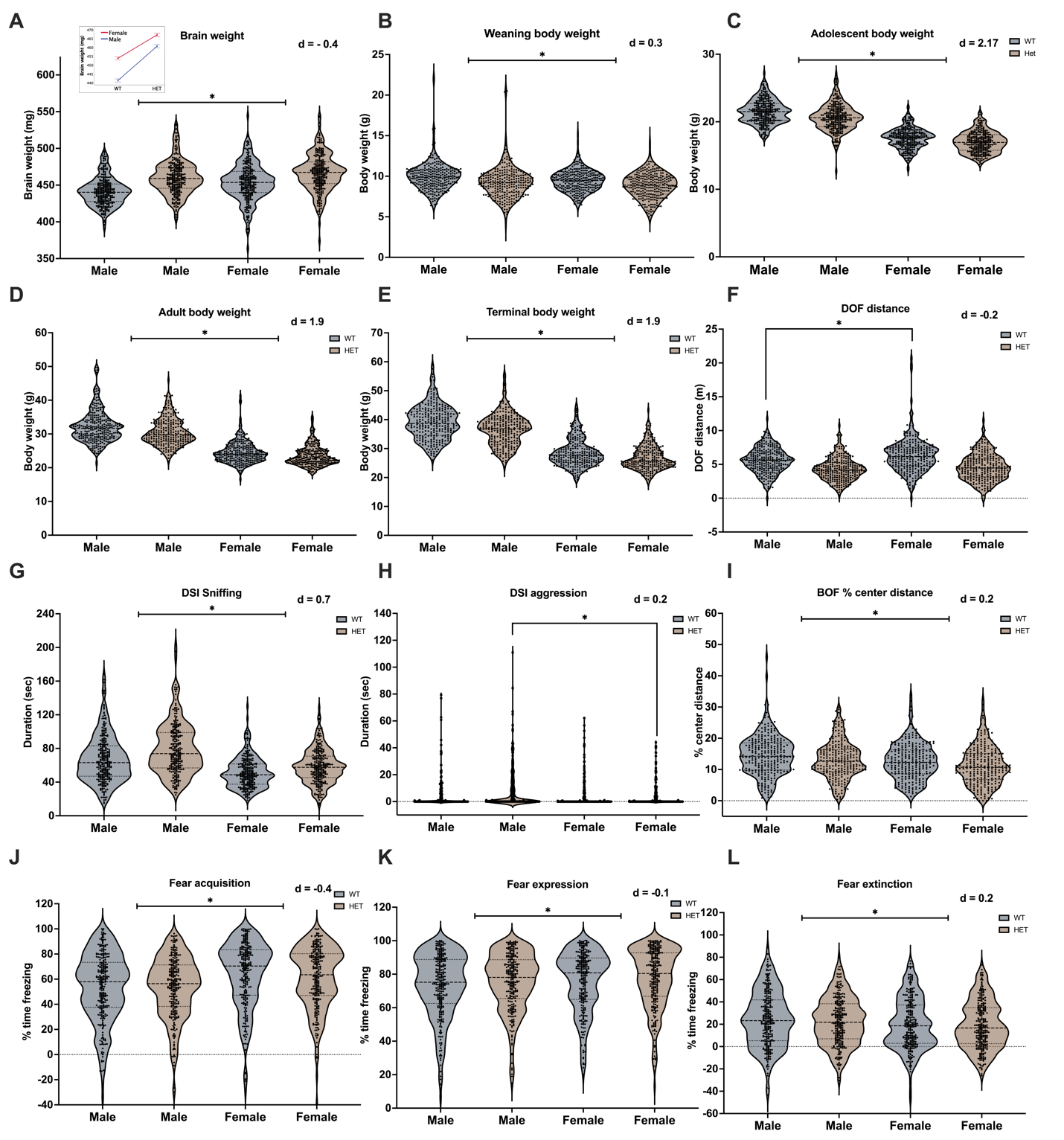
Fig. S1. Population sex differences across traits and impact of *Chd8* haploinsufficiency. Female brains weighed more than male brains (A; F_1, 1,031_ = 107.003, p < .001). A *Chd8* genotype by sex interaction revealed that males exhibited a larger effect than females on brain weight (graph insert in A; F_1, 1,031_ = 11.235, p < .001, note that graphs in A do not start at 0). Males had larger body weights than females at weaning (B; F_1, 1,041_ = 25.843, p < .001), adolescence (C; F_1, 1,039_ = 2,318.135, p < .001), young adulthood (D; F_1, 1,040_ = 1,934.43, p < .001), and end of life (E; F_1, 1,021_ = 2,313.611, p < .001). A *Chd8* genotype by sex interaction revealed that WT females traveled farther than WT males in the DOF test (F; F_1, 1,037_ = 4.289, p < .001). Males spent more time sniffing a same sex juvenile in the direct social interaction task (G; F_1, 1,035_ = 172.234, p < .001) compared to females. *Chd8* heterozygous (*Chd8^+/-^* ) males had higher DSI aggression durations compared to *Chd8^+/-^* females (H; F_1, 1,035_ = 5.848, p < .001). Percent center distance traveled in the bright open field test was higher in males than females (I; F_1, 1,038_ = 16.983, p < .001). Fear acquisition (F_1, 1,031_ = 37.289, p < .001) and fear expression (F_1, 1,036_ = 6.997, p < .01) were higher in females compared to males (J, K) while fear extinction was higher in males than females (L; F_1, 1,036_ = 10.550, p < .01). *P < .05; d = Cohen’s D effect size estimates based on the differences between the male and female populations. Negative Cohen’s D values reflect an increase in the dependent variable in females compared to males.
